# Geometric experience sculpts the development and dynamics of hippocampal sequential cell assemblies

**DOI:** 10.1101/2023.12.04.570026

**Authors:** Usman Farooq, George Dragoi

## Abstract

Euclidean space is the fabric of the world we live in. Whether and how geometric experience shapes our spatial-temporal representations of the world remained unknown. We deprived rats of experience with crucial features of Euclidean geometry by rearing them inside translucent spheres, and compared activity of large hippocampal neuronal ensembles during navigation and sleep with that of cuboid cage-reared controls. Sphere-rearing from birth permitted emergence of accurate neuronal ensemble spatial codes and preconfigured and plastic time-compressed neuronal sequences. However, sphere-rearing led to diminished individual place cell tuning, similar neuronal mapping of different track ends/corners, and impaired neuronal pattern separation and plasticity of multiple linear track experiences, partly driven by reduced preconfigured network repertoires. Subsequent experience with multiple linear environments over four days largely reversed these effects, substantiating the role of geometric experience on hippocampal neural development. Thus, early-life experience with Euclidean geometry enriches the hippocampal repertoire of preconfigured neuronal patterns selected toward unique representation and discrimination of multiple linear environments.

## Introduction

Spontaneous neuronal activity of the hippocampus embodies an internal model of external space (*1–5*), which could support its critical role in navigation (*2, 6*) and contextual learning (*7, 8*). The hippocampal network of immature experimentally-naïve rats spontaneously expresses time-compressed preconfigured preplay sequence motifs during sleep (*9, 10*) that are recruited to encode future de-novo sequential experiences on linear tracks as early as postnatal day 17 (P17) and future discrete spatial locations as early as P15 (*11*). This suggests that the internal model of linear space is largely innate (*12*) and could emerge likely via genetically-encoded developmental programs (*13*) in the absence of explicit navigational experience once synaptic wiring matures (*11, 14*). However, owing to the geometry of the spatial environment in which they were raised (i.e., a cuboid-shaped home cage), these rats were inherently exposed to linear boundaries, vertical borders, planar floors, and right-angled corners, which are crucial features of Euclidean space (Fig. 1A top). These and other properties of Euclidean space facilitate sequentialization, segmentation, and orthogonalization of spatial experiences providing direct experience with *geometric linearity within Euclidean space*, which we refer to as ‘geometric linearity’. This daily ‘unaccounted’ exposure to geometric linearity during early life may have enabled the formation of an internal model for linear space in lieu of or in addition to intrinsic developmental factors and non-spatial experiences as a form of latent learning (*6*). To disentangle the contributions of early-life latent learning in Euclidean space from those of age-related intrinsic developmental programs and non-spatial experiences to the emergence of an internal, preconfigured model of space, we deprived rats of experience with geometric linearity.

**Fig. 1.**
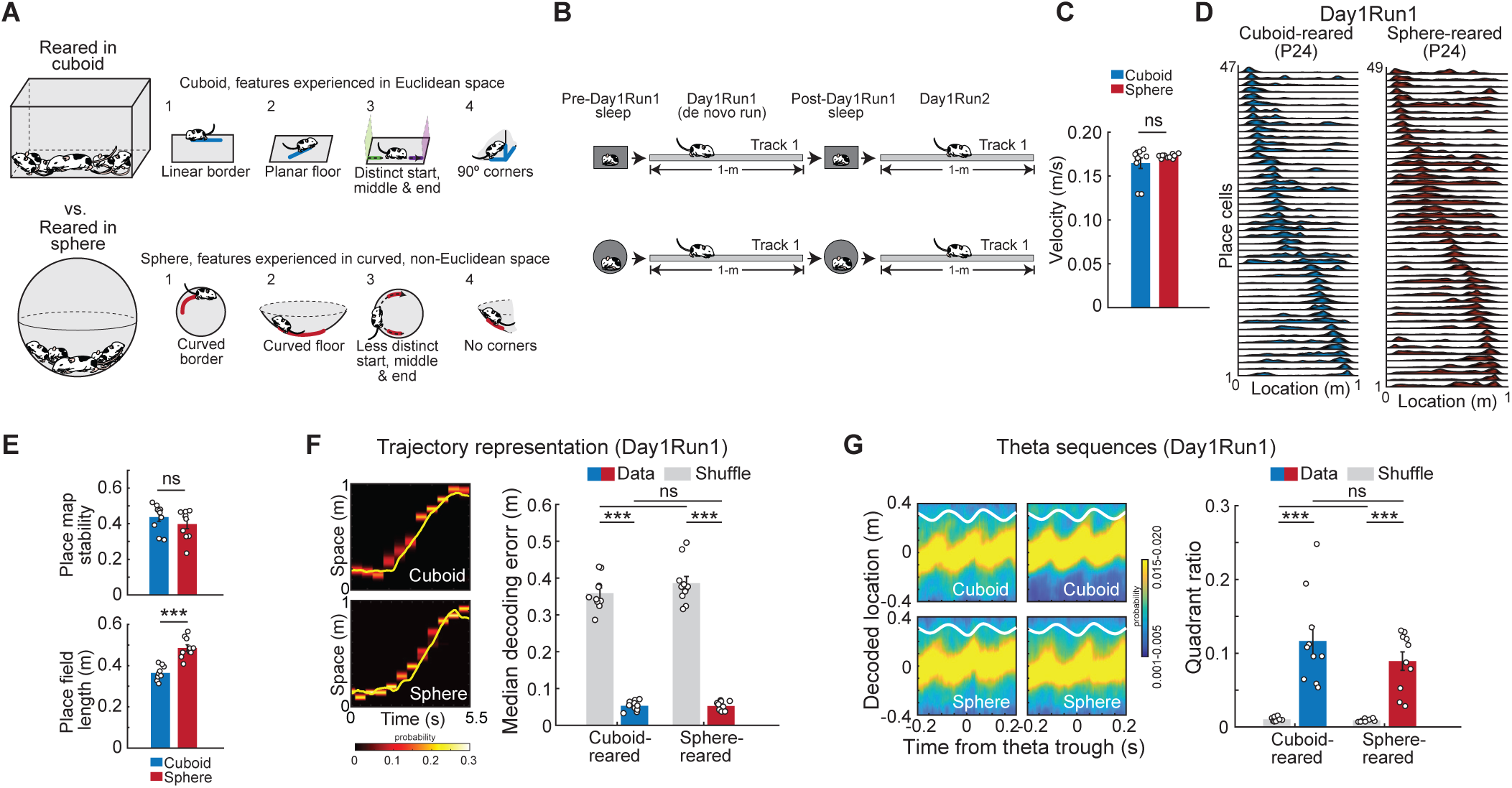
Experimental design and early-life emergence of place cells and theta sequences after sphere-rearing. (**A**) Experimental design: rearing in cuboid (top) or spherical (bottom) cages (left panel), and the critical features of Euclidean geometry experienced in cuboid-cages (blue lines, A1-4, top) and their curved non-Euclidean analogs during deprivation from Euclidean geometry inside spheres (red lines, A1-4, bottom). (**B**) Experimental design (Day1): Sleep and navigational run sessions on a linear track. (**C**) Animal velocity during run (p>0.05, rank-sum test). (**D**) Examples of simultaneously-recorded place cell sequences in cuboid-(left) and sphere-reared rats (right). (**E**) Place map stability (top, p>0.05, rank-sum test) and primary place field length (bottom, p<0.0006, rank-sum test) during Day1Run1 across experimental groups. (**F**) Decoded (heatmaps) and real (yellow lines) single-lap Day1Run1 trajectories (left). Decoding error (right) across groups (p>0.05, rank-sum tests) and compared to shuffles (p<10^-3^, signed-rank tests). (**G**) Theta sequences (left, examples) compared across groups (p>0.05, t-test) and against shuffles (p<10^-3^, paired t-tests) using the quadrant ratio method (right). N=10 (5 rats/group; 2 directions/rat). ***p<0.005. ns=not significant.

The deprived rats were raised from birth together with their littermates and the dam inside 23-inch diameter bedded, externally lighted, translucent white plastic spheres, which is one model of non-Euclidean geometry with positive curvature. The spherical environment deprived the rats of experience with: 1) linear borders and linear trajectories along them, 2) flat surfaces which permit linear trajectories, 3) right-angled corners and vertical geometric landmarks that orthogonalize and segment linear trajectories in the horizontal plane (Fig. 1A, top). Instead, the spherical home cage provided experience with continuously curved borders and concave surfaces (Fig. 1A, bottom). Considering that linear experience-related neuronal sequences critical for navigation and memory formation, adult-like hippocampal theta sequences (*15*) and plasticity in replay (*16–19*) emerge in concert at P23-24 (*11*), we deprived our rats of experience with geometric linearity until P24. We recorded the activity of large ensembles of dorsal CA1 neurons as experimentally-naïve P24-27 sphere-reared rats slept in spheres and ran on multiple linear tracks (*10*) over four consecutive days. We compared their activity across various brain states (active run behavior, awake rest and sleep) and at various levels of network organization with that of age-matched younger, and adult cuboid cage-reared control rats. Finally, our multi-day neuronal ensemble recordings from the same groups of rats allowed us to investigate whether experience of geometric linearity in the form of multiple linear tracks gathered over several days could reshape the hippocampal internal model of space in the sphere-reared rats.

## Results

### Representation of de-novo linear trajectories during run by hippocampal neuronal ensembles in the sphere-reared rats

Sphere– and cuboid-reared rats were implanted bilaterally with Neuronexus silicon probes (total of 64-128 recording sites) over dorsal CA1 area of the hippocampus (*11*) at P20-21 and returned to their respective home cages. Over the next 2-3 days, the electrodes were gradually moved toward and into the CA1 pyramidal layer while the rats were resting and sleeping in small, heated sleep boxes of corresponding geometry that continued to deprive the sphere-reared rats of physical and visual exposure to geometric linearity. Activity from CA1 neuronal ensembles and local field potentials were recorded (Fig. 1B) at P23-24 on Day1 while the naïve animals sequentially: 1) slept in the box for ∼90 min (Pre-Day1Run1 sleep), 2) explored for the first time a 1 m-long linear track in the same room for ∼30 min (de-novo Day1Run1), 3) slept in the box for ∼90 min (Post-Day1Run1 sleep), and 4) ran on the same track for ∼30 min (Day1Run2). Sphere-rearing did not significantly influence locomotor dynamics during development, as evidenced by similar individual rat locomotion when co-housed with the littermates and mother in rearing contexts of corresponding geometry, as tested at different developmental ages (N=7 cuboid-reared and N=8 sphere-reared rats) using multi-animal DeepLabCut software to estimate rats’ positions (*20*). Furthermore, the geometry of the rearing environment did not influence the rats’ velocity and overall behavior on linear tracks (Fig. 1C). In addition, the sphere-rearing protocol did not increase stress levels in the rats compared to cuboid-rearing. This was assessed by quantifying fecal corticosterone (FC) levels of individual rats that provided a temporally-integrated non-invasive measure of chronic stress (*21*) across groups of cuboid– and sphere-reared rats exposed to linear tracks (using FC enzyme-linked immunosorbent assay ELISA: cuboid vs. sphere, 54.0±7.5 vs. 44.4±3.5 ng/g, p>0.05, t-test, N=10 rats/group). Therefore, sphere-rearing was a non-stressful protocol to probe the influence of early-life experience (or lack thereof) with geometric linearity on neurodevelopment without impacting overall locomotor dynamics.

We next recorded simultaneously from an average of 63.0±9.9 (range: 44-99 well-isolated stable CA1 neurons) and 67.6±14.3 neurons (range: 31-111 well-isolated, stable CA1 neurons) across all sleep and run sessions in P23-24 cuboid-reared (N=5, one P23 and four P24 rats) and P24 sphere-reared (N=5) rats, respectively (Fig. 1D). Place cells were expressed in the sphere-reared rats during their first-ever, de novo run session on a linear track and maintained a similar degree of place field stability (Fig. 1E, top). This indicates that sphere-rearing did not prevent these rats from rapidly expressing a stable representation of the linear space at P24, despite lacking prior experience of this sort. The rapid stable expression of place maps is in contrast with the gradual age-dependent postnatal development of these features in younger P15-P22 cuboid-reared rats (*11*). However, sphere-reared P24 rats displayed larger primary place field lengths for linear space, a metric for spatial tuning, compared with P23-24 cuboid-reared rats (Fig. 1E, bottom), likely contributed by their lack of early-life experience with critical features of linear geometry (Fig. 1A). Despite this difference in individual place cell tuning, animals’ linear trajectories were represented with similarly high precision by ensembles of place cells across both experimental groups using Bayesian decoding at the behavioral timescale (*10, 11, 22*) (Fig. 1F; cuboid vs. sphere: 5.3±0.4 vs. 5.2±0.4 cm trajectory decoding median error, p>0.05, rank-sum test).

Hippocampal theta sequences, which bind recent-past, present, and immediate-future visited track locations in temporally-compressed manner within individual cycles of theta oscillation during locomotion (*15, 23*), emerge at P23-24 in control cuboid-reared rats (*11*). In P24 sphere-reared rats, we observed similar theta oscillation frequency during run (6.7±0.2 vs. 6.8±0.1, cuboid vs. sphere-reared, p>0.05, rank-sum test) and similar theta sequence compression and spatial binding within theta oscillations while rats traversed the middle portion of the track (theta sequence quadrant ratio: 0.12±0.02 vs. 0.09±0.01, cuboid vs. sphere-reared, p>0.05, t-test; Fig. 1G). Our results indicate that the emergence of hippocampal neuronal ensembles’ competence to accurately represent animal position and linear trajectory at both behavioral timescale and temporally-compressed theta scale during de novo run (*11, 15*) does not require prior experience with geometric linearity.

### Depiction of de-novo linear trajectories during sleep and awake rest by hippocampal neuronal ensembles in the sphere-reared rats

During sleep and rest, neuronal sequences replay experienced (*19, 24, 25*) and preplay future novel (*9, 10*) extended linear animal trajectories in a time-compressed manner. These p/replay events occur within brief epochs (100-800 ms) of elevated population neuronal activity often associated with ripple oscillations called frames (*10*). P23-24 cuboid-reared animals naïve to extended linear space exhibited significant preplay and replay of the de-novo Day1Run1 trajectory during sleep and rest frames (Fig. 2A) and experience-related plasticity expressed as stronger trajectory depiction in replay compared with preplay during sleep (*11*) (Fig. 2C). The proportions of frames depicting experience-related trajectories occurred at similar levels to previously reported p/replay in adult rats (*10, 11, 16, 26, 27*). Despite being deprived of prior experience with geometric linearity, P24 sphere-reared animals displayed robust extended linear preplay during Pre-Day1Run1 sleep and replay in Post-Day1Run1 sleep (Fig. 2, B and C). This demonstrates that the emergence of network preconfiguration into temporally-compressed neuronal sequences that preplay a future novel extended linear trajectory in experimentally-naïve animals does not require prior experience with geometric linearity. Whereas sphere-reared animals had lower incidences of preplay and replay compared with their cuboid-reared counterparts, these levels of preplay were sufficient to support experience-related plasticity in neuronal sequence replay (*11, 16, 18*) during Post-Day1Run1 sleep (Fig. 2C). These results indicate that network competence to rapidly encode and store a sequential linear spatial experience emerges in the hippocampus despite the lack of prior experience with geometric linearity, likely aided by the experience-independent emergence of preplay.

**Fig. 2.**
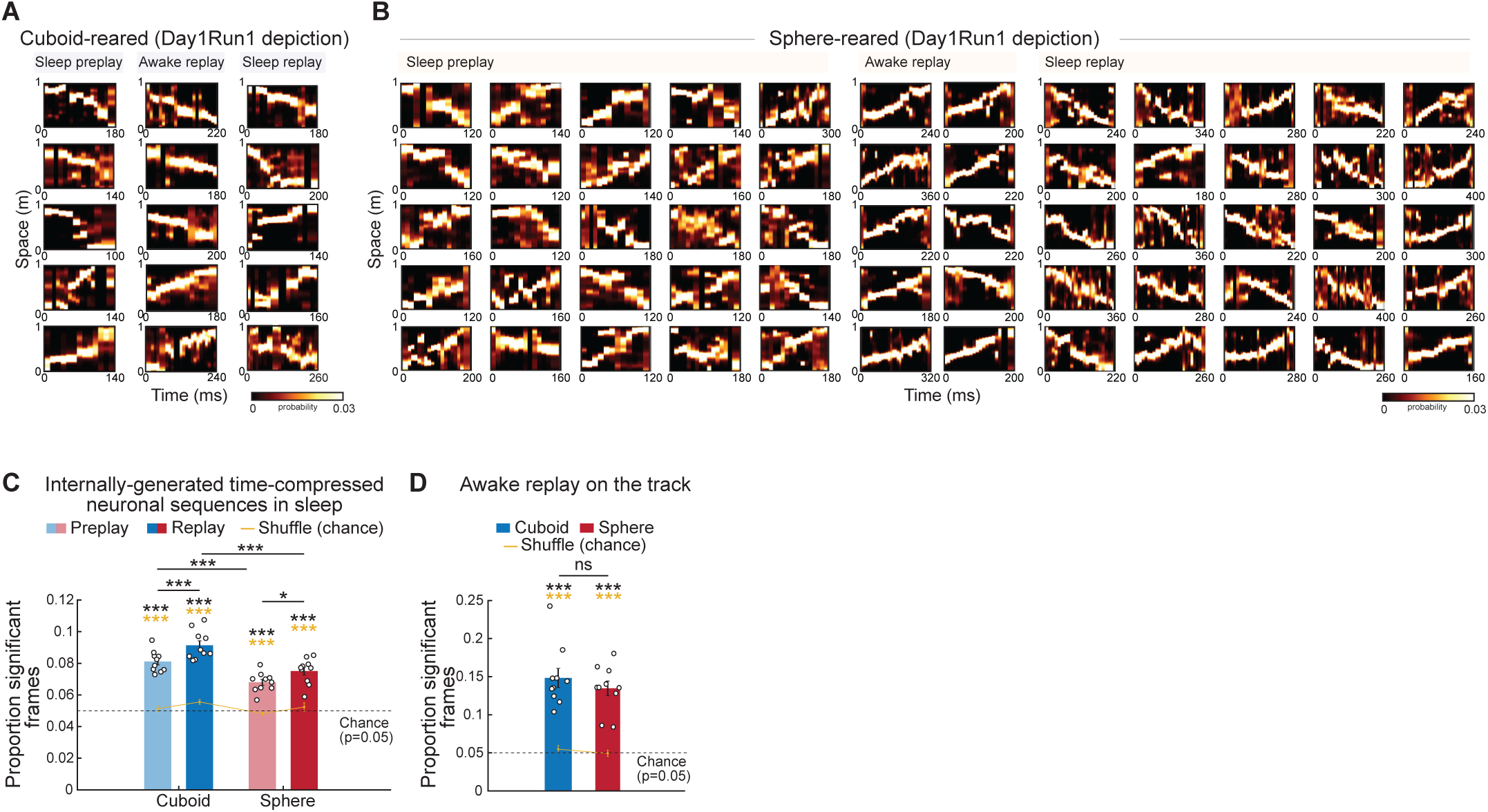
Early-life emergence of sleep preplay, awake replay and plasticity in sleep replay after sphere-rearing. (**A-B**) Examples of Day1: sleep preplay, on-track awake replay, and plasticity in sleep replay depicting Day1Run1 trajectories in cuboid-(A) and sphere-reared rats (B). (**C**) Significant incidence of sleep p/replay depicting Day1Run1 in both groups (cuboid and sphere p/replay: vs. shuffle: p<10^-3^, paired t-tests, yellow stars; p<10^-5^ vs. p=0.05 chance, t-tests, black stars; p<10^-10^, Binomial tests vs. p=0.05 chance for all groups and sleeps). Reduced incidence of p/replay in sphere-reared rats (p<0.0006, t-tests). Plasticity in replay (replay>preplay) in both groups (p<0.03, paired t-tests). (**D**) Incidence of on-track awake replay (cuboid and sphere p/replay: vs. shuffle: p<10^-4^, paired t-tests, yellow stars; vs. p=0.05 chance: p<10^-4^, t-tests, black stars; p<10^-10^, Binomial tests vs. p=0.05 chance for both groups) is similar across groups (p>0.05, t-test). N=10 (5 rats/group; 2 directions/rat). ***p<0.005. *p<0.05. ns=not significant.

Awake-rest replay, time-compressed representation of experienced trajectories during immobility on the track (*24*), has been implicated in spatial navigation planning and stabilization of cognitive maps. Awake-rest replay occurred with similar incidence in the cuboid– and sphere-reared rats during the immobility epochs of the de novo Day1Run1 (Fig. 2, A to B and D).

Together, these results indicate that while prior experience with linear geometry was dispensable for the emergence of neuronal sequences depicting extended linear trajectories during sleep, it nevertheless contributed towards the incidence of trajectory-depicting sequence p/replay in sleep. This is suggestive of an influence of prior latent geometric experience toward (pre)configuration of the hippocampal network dynamics recruited during future novel experiences.

### Similarity in hippocampal ensemble depiction of distinct linear track ends and corners altered representation of linear space after sphere-rearing

Sphere-reared rats were deprived from birth of experience with geometric linearity, including experience of environmental ends/corners and vertical borders and, instead, accrued excess experience with curved space during early postnatal neuro-development, which attenuated the incidence of time-compressed linear p/replay (Fig. 2). We asked whether these differences in geometric experience and preplay levels due to sphere-rearing also affected the network ability, at any level, to reliably map individual locations along an extended linear space. Therefore, we computed the similarity between the neuronal ensembles mapping pairs of locations along the linear track during Day1Run1. In cuboid-reared rats, locations near the two track ends (points furthest apart on the linear track) recruited distinct neuronal ensembles that exhibited low population vector correlations (Fig. 3, A and B). In contrast, in sphere-reared rats, pairs of locations situated near the opposite track-ends recruited more similar place cell ensembles at P24 compared with cuboid-reared rats (Fig. 3B).

**Fig. 3.**
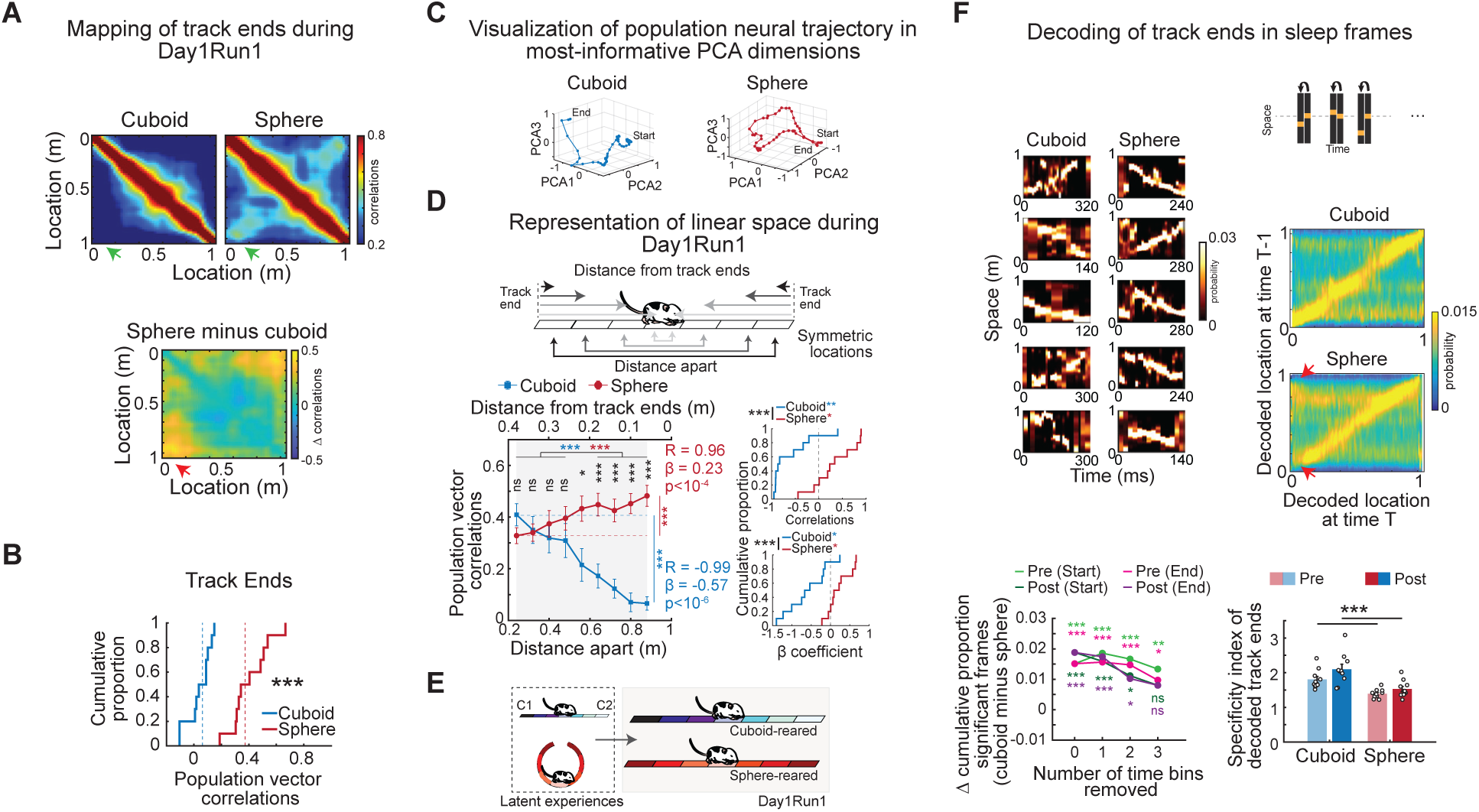
Changes in hippocampal ensemble depiction of linear track ends/corners after sphere-rearing. (A) Average correlation matrix of neuronal ensemble mapping of locations along the track during Day1Run1 for all cuboid-(top-left), sphere-reared rats (top-right), and the difference between groups (bottom). Green/red arrows mark higher similarity of population vectors representing track ends in the sphere-reared rats. (**B**) Population vector correlation of Day1Run1 place maps across different segments of the track. Note higher correlations between track ends (p<0.0002) in sphere-vs. cuboid-reared rats (rank-sum tests). Vertical dotted lines mark corresponding median values. (**C**) Visualization of the neural manifold depicting a 1-m-long linear track during Day1Run1 across the first 3 most informative dimensions of the principal component space. Note a more curved neuronal representation of space in a sphere-, but not cuboid-reared rat. (**D**) Schematic (top) and values (bottom-left) of population vector correlations between place maps at locations equidistant/symmetric from the two ends of the track (and track middle) as a function of distance between the corresponding locations during Day1Runs. Grey mask: range of locations for which a Pearson’s correlation (R) and least-squares linear regression line was computed between the group average population vector correlations of place maps and the corresponding linear distances separating the locations of those place maps. Stars indicate significant differences at specific distances in cuboid-vs. sphere-reared rats (rank-sum tests). Dashed red and blue lines depicts average value for sphere– and cuboid-reared rats at 0.24 m distance apart to aid comparison. Significant decrease and increase in population vector correlations in cuboid– and sphere-reared rats as distance increases (closest vs. furthest location – 0.24 vs. 0.88 – cuboid: p<10^-4^, sphere: p<0.004, signed-rank tests, depicted by stars on right of plot; four spatial locations in middle vs. four spatial locations at ends, cuboid: p<10^-9^, sphere: p<10^-3^, rank-sum tests, depicted towards top of plot). Bottom-right: Correlations (top) and beta-coefficients (ß, slope) of the best-fit linear regression line (bottom) for all individual animals and run directions in cuboid– and sphere-reared rats (cuboid-vs. sphere-reared, p<0.003, rank-sum tests; cuboid: correlations<0, p<0.01 and beta-coefficients<0, p<0.02, sphere: correlations>0, p<0.05 and beta-coefficients>0, p<0.05, signed-rank tests, colored stars). (**E**) Color-coded cartoon depicting increased similarity of neuronal ensembles active at track ends/corners (represented by more similar colors) and its gradual decrease between symmetric track locations as a function of distance from ends in sphere-reared rats during Day1Runs. We propose this effect reflects early latent experiences in cuboidal or spherical home cages (depicted by a line or a ring respectively). C1, C2 = corner1-2. (**F**) Left-top two columns: Examples of Day1Run1 trajectory decoding during Post-Day1Run1 sleep frames in cuboid (left column) and sphere-reared rats (right column) illustrating confusion of track ends after sphere-rearing. Note relative swapping of decoding probabilities for the two track ends in these examples from sphere-reared rats. Left-bottom: Increased similarity in the cumulative proportions of significant frames between the cuboid and sphere-reared rats in Pre-Day1Run1 and Post-Day1Run1 sleep as a function of the number of 20 ms-long bins removed from the beginning or end of the originally non-significant sleep frames. Similarity was assessed via Z-tests for 2 proportions on pooled data across all animals for Post– and Pre-Day1Run1 sleep. Right-top: Schematic of the method for computing time-space contingency analysis. Right-center: Average time-space contingency matrices of decoded locations along the linear track; arrowheads mark swapping of decoding of track ends in sphere-, but not cuboid-reared rats. Right-bottom: Specificity index of decoded track ends (p<0.003, t-tests). N=10 (5 rats/group; 2 directions/rat). ***p<0.005. **p<0.01. *p<0.05. ns=not significant.

We next investigated whether the representation of continuous linear space was affected beyond the observed similarity of track ends’ representation after sphere-rearing. To understand the structure of population activity recruited by running on a 1 m-long linear track, we first visualized the neural ‘trajectory’ taken by the population of location-normalized place cell ensembles in the top 3 dimensions of the principal component neuronal space. Visual inspection revealed a higher curvature in the neural trajectory of the sphere-reared compared with cuboid-reared rats (Fig. 3C). To quantify this phenomenon, we compared population vector correlations between place maps in the non-reduced data at locations equidistant on the linear track using the two track ends as our references (these locations were also symmetric by the track middle, as an alternate reference frame) (Fig. 3D, top). In P23-24 cuboid-reared animals, the neuronal representations of locations situated further apart in linear space were more distinct than those of nearby locations (Pearson’s correlation for mean values across all rats, group means R: –0.99, slope of regression fit (β): – 0.57, p<10^-6^; Fig. 3D, bottom-left). This network property allowed for mapping and discrimination of distinct locations along a linear space. In contrast, sphere-reared rats displayed a significant positive relationship between the distance separating equidistant locations on the track and the similarity of recruited neuronal ensembles (Fig. 3D, bottom and right; group average and individual rats presented, respectively). Thus, due to the higher similarity of track ends, the remaining locations on the track as the animal approached the track ends, computed as distance-to-closest-end (*28*), also became increasingly similar in the sphere-reared rats, resulting in an altered representation of the linear space. A downstream neuronal reader might interpret these altered dynamics in representing linear space in sphere-reared rats as a deviation from linearity, or ‘warping’. Notably, despite signatures of a different mapping of linear space by a subgroup of neurons in the sphere-reared rats, overall, sequential track locations were represented sufficiently differently to enable an accurate representation of animals’ sequential trajectories. Thus, despite the lack of any prior experience with geometric linearity, preconfigured sequential motifs were rapidly recruited to accurately represent de novo linear experiences in the hippocampus of sphere-reared rats (Fig. 1F), suggesting that the dominant neuronal representation at the behavioral timescale in the hippocampus is linear in nature.

Based on these findings we propose that: 1) early-life experience with geometric linearity, start and end vertical landmarks, and right-angled corners, which segment linear boundaries in cuboid, but not spherical environments, shaped the hippocampal network to discriminate and respond more specifically to these orienting landmarks later in life, and 2) early-life experience with curved space induced an altered representation of the external linear space in the hippocampus (see cartoon model in Fig. 3E).

Given that novel place cell sequences for linear space are p/replayed during the preceding/following sleep (*5, 9, 11, 19*), we next investigated the effect of increased experience with non-linear, curved space on the decoded trajectories during sleep. We wondered if the changed representation of linear space in the sphere-reared rats could be related to a lower incidence of linear preplay sequences compared to cuboid-reared rats (Fig. 2C). To investigate if the signature of these changes were observed in sleep frames, we focused on the depiction of the two track ends given that they were more likely to recruit similar ensembles during run (Fig. 3B). We observed a network ‘confusion’ of the two track ends in a small subgroup of decoded frames in the sphere-reared rats (selected example frames in Fig. 3F, left). This confusion was primarily expressed as a co-representation or ‘swapping’ of the two track ends along with an accurate depiction of the remaining part of the decoded trajectory along the track during sleep.

We designed two separate analyses to quantify this altered depiction of linear space by neuronal sequences during time-compressed sleep frames in the sphere-reared rats. First, since the swapping/confusion of track ends was most apparent in the start or end of sleep frames, we generated surrogate frames with gradually-longer 20-60 ms-long epochs (1-3 time bins, 20 ms/bin) removed from either the start or the end of the initially non-significant frames, called reduced-frames. We quantified the sequential trajectory content depicted in the remaining part of the reduced-frame by comparing its weighted correlation with its time bin shuffles (as in Fig. 2). With each iteration of 20 ms epoch removal, we quantified the proportion of additional reduced-frames that became significant for decoded trajectories in sleep frames after bin(s) removal in cuboid– and sphere-reared rats. Removal of three 20 ms-long epochs from either the start or end of the sleep frames was sufficient to reduce the difference between the cumulative proportion of significant frames of cuboid– and sphere-reared rats in Pre– and in Post-Day1Run1 sleep sessions (Fig. 3F, bottom-left).

Second, we developed a method that uncovered the short timescale average spatial relationships represented within frames. We constructed a contingency matrix that displayed the relationship of relative spatial locations represented across consecutive 20 ms temporal bins in sleep frames, a measurement of the degree of similarity of representation of locations in consecutive time bins. In cuboid-reared rats, both in Pre– and Post-Day1Run1 sleeps, we discovered a robust average likelihood to depict similar/nearby locations in consecutive time bins within frames. This was impaired in the sphere-reared rats specifically on or near track ends and manifested as a decrease in specificity of depiction of the two track ends in the sphere-reared compared with cuboid-reared rats (Fig. 3F, top-right). We computed a specificity index for the track ends, which was significantly lower in the sphere-compared with cuboid-reared rats in both the Pre– and Post-Day1Run1 sleep (Fig. 3F, bottom-right). Indeed, for sphere-, but not cuboid-reared rats, there was a significant correlation between the proportion of significant sleep p/replay frames (Fig. 2C) and the sleep specificity index for the decoded track ends (cuboid: R = –0.01, p>0.05; sphere: R = 0.49, p<0.03, Spearman’s Rank-order correlations). Meanwhile, no relationship was observed between the proportion of significant sleep p/replay frames and the average spatial information during run (cuboid: R = –0.04, p>0.05; sphere: R = 0.28, p>0.05, Spearman’s rank-order correlations).

These results indicate that the changes in representation of linear space observed in the sphere-reared rats during run (Fig. 3, A to D) were accompanied by similar changes in spatial-temporal dynamics during sleep (Fig. 3F). Overall, these changes following sphere-rearing are suggestive of a degree of representational ‘warping’ of linear space in these rats due to a lack of experience with geometric linearity and track ends/corners that was replaced by an increased experience with curved space during postnatal brain development.

### Reduced repertoire of clustered neuronal ensembles during sleep after sphere-rearing

Given the increased likelihood of experience with orthogonalization and segmentation of trajectories in cuboid-vs. sphere-reared reared rats due to presence of 90-degree corners and linear vertical borders (Fig. 1), we wondered if these different experiences across development shaped animals’ internal models of space accordingly. To test this hypothesis, we investigated the structure of spontaneous activity of hippocampal neuronal ensembles during sleep (before animals’ first exposure to an extended linear track), which was likely driven by the intrinsic dynamics of the network while inputs from the external world are minimally processed. We discovered that neuronal activity of pairs of putative pyramidal cells during Pre-Day1Run1 sleep (defined for each neuron as their spike counts in each of all the sleep frames) exhibited significantly higher correlations in the sphere-reared compared with cuboid-reared rats (Fig. 4, A and B). This difference between groups remained significant after accounting for possible contributions from changes in firing rates (*29*) by normalizing the correlation values (expressed as values ranging from 0 to 1) using firing rate-matched shuffles (Fig. 4C). A higher proportion of cell pairs were significantly more correlated with each other than their firing-rate matched shuffles during sleep in sphere-vs. cuboid-reared rats (34% vs. 15% at a cutoff p-value of 0.05, Fig. 4C inset). These results suggest that altered early-life geometric experience due to sphere-rearing impacted neuronal pairwise relationships of a significant proportion of the hippocampal neuronal network active during sleep.

**Fig. 4.**
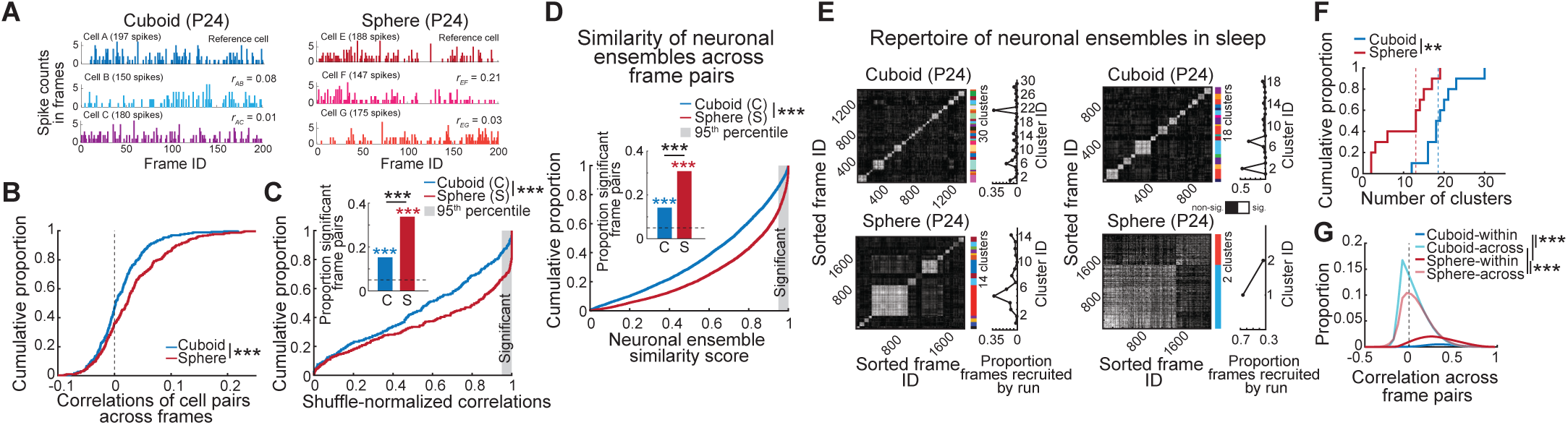
Reduced repertoire of distinct neuronal ensembles during sleep after sphere-rearing. (**A**) Examples of spike count correlations between cell pairs across 200 frames for 3 pyramidal cells during Pre-Day1Run1 sleep in cuboid-(left) and sphere-reared (right) rats. Top: reference cell for correlations. Total spike counts are given in brackets next to cell ID. (**B**) Cumulative distributions of correlations of spike counts between pyramidal cell pairs across sleep frames. Note higher values in sphere-vs. cuboid-reared rats (p<10^-4^, rank-sum test). (**C**) Shuffle-normalized correlations of cell pairs are higher in sphere-vs. cuboid-reared rats (p<10^-5^, rank-sum test). Inset: proportion of significantly correlated cell pairs (>95^th^ percentile of shuffles) is higher in sphere-vs. cuboid-reared rats (p<10^-7^, Z-test for 2 proportions). (**D**) Similarity scores (p<10^-10^, rank-sum test) and proportions of correlated Pre-Day1Run1 sleep frame pairs (inset; p<10^-10^, Binomial tests) vs. chance and between experimental groups. Note higher scores and proportions in sphere-reared vs. control rats (p<10^-10^, Z-test for 2 proportions). (**E**) Examples of clustered Pre-Day1Run1 sleep frames with correlated activity in cuboid-(top) and sphere-reared (bottom) rats. Colorbar: color-coded clusters and cluster demarcation. Curves to the right of cluster panels: Proportions of frames from each cluster that are significantly correlated with the future run trajectory. (**F**) Number of clusters during Day1 sleep sessions (cuboid>sphere, p<0.006, rank-sum test). N=10 (5 rats/group; 2 sleep sessions/rat). (**G**) Distributions of Pre-Day1Run1 sleep frame-pair correlations (within>across clusters, p<10^-10^; sphere vs. cuboid, within or across clusters: p<10^-10^, exact permutation tests). (B-D, G) N=5 rats/group. ***p<0.005. **p<0.01.

The distinctiveness of neuronal ensembles activated across sleep frames provides a readout of the intrinsic p/replay repertoire for unique depiction of multiple future and past experiences (*10*). We investigated whether the increase in cell-pair correlations during sleep after sphere-rearing (Fig. 4, A to C) translated into an increased similarity in neuronal ensembles activated across frames. Thus, for each sleep frame we generated a population vector composed of the spike counts (≥0) emitted in the frame by each of all recorded putative pyramidal neurons. For each of all frame pairs, we derived a neuronal ensemble similarity score (i.e., the shuffle-normalized correlation expressed as a value ranging from 0 to 1) between the corresponding pair of neuronal population vectors. In agreement with our cell-pair correlation results (Fig. 4, A to C), we found that neuronal ensembles activated across pairs of sleep frames were overall more similar in naïve sphere-reared compared with cuboid-reared rats (Fig. 4D). Groups of sleep frames sharing similar neuronal ensembles formed distinct clusters (*30*) (detected using k-means clustering algorithm; Fig. 4E). Importantly, cuboid-reared rats had a larger repertoire of such clusters compared to sphere-reared rats (cuboid vs. sphere median number of clusters: 18.5 vs. 13 in Day1 sleep sessions, p<0.006, rank-sum test, Fig. 4F, which differed from shuffled datasets in both groups). The within-cluster correlations were higher than the across-cluster ones (within>across clusters in both rat groups, p<10^-10^; sphere vs. cuboid, within clusters or across clusters: p<10^-10^, exact permutation tests, Fig. 4G). In addition to the fewer clusters of correlated frames, sphere-rearing led to reduced intra-cluster and increased between-cluster similarities in neuronal ensemble activity across frame-pairs compared with cuboid-reared rats (Fig. 4G). Neuronal ensembles active within a subset of sleep clusters were later preferentially recruited to represent future Day1Run1 experiences in both experimental groups, indicating these pre-configured ensembles are indeed selected for representation of future novel experiences (*1*) (Fig. 4E, right).

Based on these findings, we propose that deprivation of critical experience with geometric linearity and exposure to geometric curvature by sphere-rearing from birth drove a reduction in cluster repertoire size.

### Impaired neuronal pattern separation and plasticity of multiple linear track experiences after sphere-rearing

As the internal model of space of sphere-reared rats during sleep relied on a reduced repertoire of orthogonal neuronal ensembles (Fig. 4), we hypothesized this could reduce the network ability to simultaneously represent multiple distinct linear track experiences by distinct neuronal ensembles. Given that sphere-reared rats were deprived of experience with right-angle corners (Fig. 1A), we exposed our rats to alternating experiences on three distinct 1 m-long linear tracks arranged in U-shape, separated by 90° corners (*10*). On Day1, rats were familiarized with running on track 1 (Figs. 1 to 4). On Day2, after a sleep session in a corresponding-geometry box (Pre-Day2Runs sleep), the rats explored track 1 in isolation followed by exploration of the contiguous 3 tracks (tracks 1-3, Day2Run2), a Post-Day2Runs sleep session in the box, and a re-exploration of the 3 tracks (Fig. 5A).

**Fig. 5.**
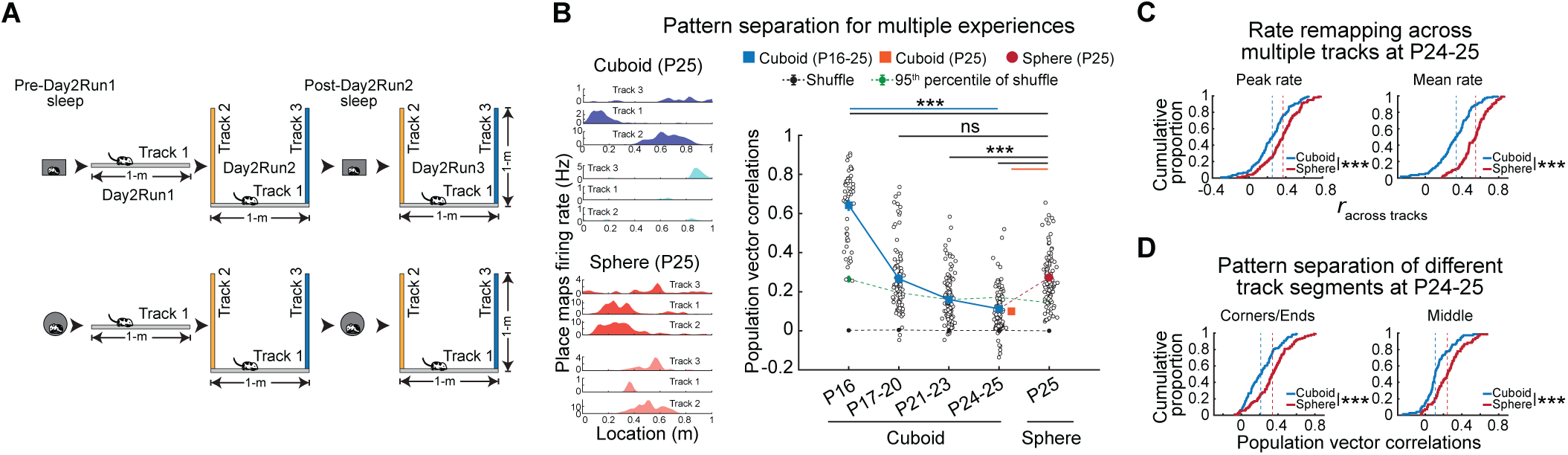
Impaired pattern separation of distinct multiple linear track experiences after sphere-rearing. (**A**) Experimental design (Day2): sleep and run on 3 linear tracks. (**B**) Developmental emergence of place cell re/mapping and pattern separation across 3 tracks. Left: examples of re/mapping in cuboid– and sphere-reared rats. Right: age-related increase in pattern-separation (re/mapping) in P16-P25 cuboid-reared rats (p<10^-10^, ANOVA) and its impairment in P25 sphere-reared rats (P25 sphere vs. P16 cuboid: p<10^-10^; P25 sphere vs. P17-20 cuboid: p>0.05; P25 sphere vs. P21-23 cuboid: p<10^-9^, P25 sphere vs. P24-25 cuboid: p<10^-10^, P25 sphere vs. P25 cuboid: p<10^-10^, rank-sum tests). N (rats/group x 2 directions/rat for each track): Cuboid: 6 (P16), 10 (P17-20), 12 (P21-23), 10 (P24-25), 8 (P25); Sphere: 10 (P25). (**C**) Reduced discriminability between multiple tracks (i.e., increased rate correlations) after sphere-rearing (P24-25 cuboid-reared vs. P25 sphere-reared: p<10^-4^, rank-sum tests). (**D**) Higher similarity between mapping of different corners/ends (p<10^-5^, rank-sum test, left) and tracks’ middle across the 3 tracks (p<10^-4^, rank-sum test, right) in the sphere-vs. cuboid-reared rats. N (rats/group x 2 directions/rat for each track): (C-D) Cuboid, P24-25 (10); Sphere, P25 (10). ***p<0.005. ns=not significant.

Since it has remained unknown when the hippocampal network first acquires the ability to distinctly represent, or pattern separate, multiple linear spatial experiences (*31, 32*) during early postnatal life, we first studied the development of pattern separation between 3 distinct linear tracks in younger, P16-25 cuboid-reared rats (N=19 rats undergoing the Day2 experimental protocol for the first time across various developmental ages). We started from P16, when time-compressed sequential motifs are not yet expressed, and continued until P25, when preplay, theta sequences, and plasticity in replay have already emerged (*11*). At younger ages, neuronal ensembles failed to discriminate (*32*) (i.e., to pattern separate) between the 3 tracks. Separation of representational patterns for the 3 tracks emerged and improved at later ages, as shown by a marked age-dependent reduction in correlations between the ensembles recruited across pairs of the 3 experiences (Fig. 5B). Interestingly, the P25 sphere-reared rats had a higher deficit in pattern separation than the P21-23 and P24-25 cuboid-reared rats (Fig. 5B), likely contributed by significantly higher correlations between population firing rate-representations of the 3 tracks (Fig. 5C). This increased place code similarity for multiple tracks in the sphere-reared rats was observed for mapping right-angle corners separating orthogonal track-pairs and track ends, and extended to other locations on the track as well (Fig. 5D).

Experience with multiple contexts over short time intervals, as in our Day2 experiment, could challenge the hippocampal network while distinctly encoding and storing these experiences due to possible memory interference (*33*). One potential solution for this problem would be the availability and selection of specific, unique preconfigured preplay patterns for encoding distinct run experiences, which could greatly reduce the interference in their representation and subsequent long-term storage (*1, 10*). Both sphere– and cuboid-reared animals at P24-25 exhibited significant preplay of the 2 novel tracks during the sleep session before they experienced the combined 3 tracks on Day2 (Fig. 6A), like previously reported in adult rats (*10*). Similar to Pre-Day1Run1 (Fig. 2C), sphere-reared rats exhibited lower incidence of preplay compared to their cuboid-reared counterparts on Pre-Day2Run2 sleep (Fig. 6A). Next, we studied if these preplay sequences uniquely depicted each of the multiple tracks by assessing the incidence of track-specific sleep preplay (i.e., frames which selectively depicted sequential trajectories for only one novel, but not the other two spatial experiences). Track-specific preplay occurred above chance levels in the cuboid-reared rats (Fig. 6A), while in the sphere-reared rats, the incidence of track-specific preplay was lower and did not exceed chance levels (Fig. 6A). This difference between rat groups indicates a deficiency in generating distinct temporal codes for multiple future experiences during sleep in the sphere-reared rats.

**Fig. 6.**
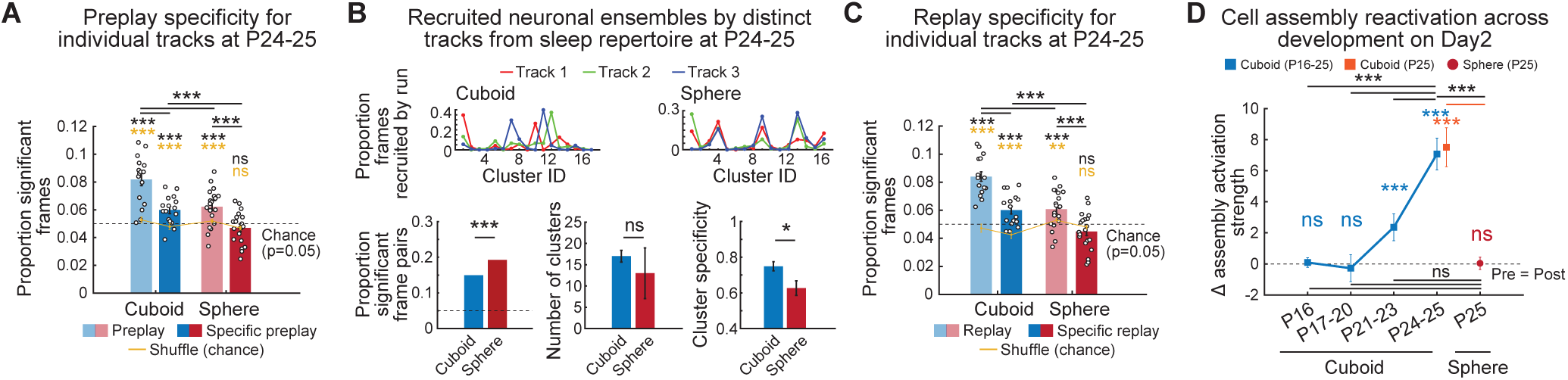
Reduced neuronal ensemble track-specific depiction of future and reactivation of past distinct multiple linear track experiences during sleep after sphere-rearing. (**A**) Proportions of preplay and ‘specific-track’ preplay after cuboid-rearing (p<10^-4^, p<0.002, paired t-tests vs. shuffle, yellow stars; p<10^-5^, p<10^-3^, t-tests vs. p=0.05 chance, black stars; p<10^-10^, p<10^-6^, Binomial tests vs. p=0.05 chance) and sphere-rearing (p<0.003, p>0.05, paired t-tests vs. shuffle, yellow stars; p<10^-3^, p>0.05, t-test vs. p=0.05 chance, black stars; p<10^-10^, p>0.05, Binomial tests vs. p=0.05 chance, respectively). Note significantly lower preplay and track-specific preplay in sphere-reared rats compared to cuboid reared rats (p<0.0007, p<0.002, t-tests). (**B**) Experiences on distinct tracks during Day2Run2 recruit neuronal ensemble from a larger repertoire of Pre-Day2Run1 sleep frame clusters in cuboid-vs. sphere-reared rats (individual rat examples are shown). The proportion of significant frame pairs in Pre-Day2Run1 sleep remain significantly higher in the sphere-vs. cuboid-reared rats (bottom-left, p<10^-10^, Z-test for 2 proportions), which does not translate to a significantly difference in the number of discrete clusters (bottom-center, p>0.05, rank-sum test). Specificity (bottom-right) of cluster recruitment from Pre-Day2Run1 sleep by each future track is higher in cuboid-vs. sphere-reared rats (p<0.03, t-test). (**C**) Proportions of replay and ‘specific-track’ replay after cuboid-rearing (p<10^-6^, p<10^-3^, paired t-tests vs. shuffle, yellow stars; p<10^-7^, p<0.002, t-tests vs. p=0.05 chance, black stars; p<10^-10^, p<10^-4^, Binomial tests vs. p=0.05 chance) and sphere-rearing (p<0.008, p>0.05, paired t-tests vs. shuffle, yellow stars; p<10^-3^, p>0.05, t-tests vs. p=0.05 chance, black stars; p<10^-10^, p>0.05, Binomial tests vs. p=0.05 chance, respectively). Note significantly lower replay and track-specific replay in sphere-reared rats compared to cuboid-reared rats (p<10^-4^, p<0.0009, t-tests) (**D**) Cell assembly plasticity (sleep reactivation exceeding pre-activation) for multiple experiences in younger cuboid-reared rats (p>0.05 for P16, P17-20; p<0.0004 for P21-23, p<10^-4^ for P24-25, and p<0.0005 for P25, signed-rank tests). Cuboid-reared P24-25 rats exhibit a multi-fold increase in experience-dependent cell assembly plasticity (sleep reactivation), which is significantly higher than in younger rats (p<0.0002, rank-sum tests). Cell assemblies in sphere-reared rats (P25) have impaired experience-related plasticity compared with P24-25 cuboid-reared rats and age matched P25 cuboid-reared rats (p<10^-5^, rank-sum tests), and similar reactivation to P16-23 cuboid-reared rats (p>0.05, rank-sum tests). N (rats/group x 2 directions/rat for each track): Cuboid: 6 (P16), 10 (P17-20), 10 (P21-23), 8 (P24-25), 6 (P25); Sphere: 10 (P25).

In addition to the low incidence of track-specific preplay, the reduced pattern separation during subsequent run in the sphere-reared rats could be due, in part, to a higher similarity between neuronal ensemble activities within frame pairs during the preceding sleep. Furthermore, a reduced repertoire of distinct neuronal ensembles during the preceding sleep on Day2 in sphere-reared rats, similar to what we observed on Day1 (Fig. 4, D to F), could also contribute to the reduction in pattern separation across the two rat groups. Indeed, the proportions of significantly correlated frames during the Pre-Day2Run2 sleep remained higher after sphere-rearing, similar to those observed during the Pre-Day1Run1 sleep (Fig. 6B). While these differences in proportions of significantly correlated frame pairs did not directly translate into significantly lower numbers of distinct clusters of neuronal ensembles on Day2 (Fig. 6B), they were associated with less-defined cluster boundaries in the sphere-reared rats. This suggests that prior experience with geometric linearity on Day 1 contributed to a partial improvement in the cluster repertoire in the sphere-reared rats, which still remained lower defined on Day2 in the sphere-compared to cuboid-reared rats. Clusters of neuronal ensembles in cuboid-reared, but not sphere-reared, rats were recruited by each of the 3 future track experiences, with minimal overlap, indicating that they contributed to the distinct encoding of multiple tracks in the cuboid-reared rats (Fig. 6B, bottom-right). These deficits observed in the sphere-reared rats extended to the experienced-trajectory replay when sphere-reared rats exhibited significant, yet fewer, replay frames compared to cuboid-reared rats, while track-specific replay was at chance level (Fig. 6C).

To investigate the experience-related network plasticity for multiple experiences, we isolated cell-assemblies active within 20 ms time bins that were significantly co-activated during the novel run experiences rats (*16, 34–36*) and quantified their activation in the sleep sessions before and after the novel experiences. We first studied the effects that impaired pattern separation in younger cuboid-reared rats (P16-22, Fig. 5B) had on cell assembly plasticity for multiple, novel experiences (Fig. 6D), as this relationship had remained unexplored. Younger cuboid-reared rats lacked, or exhibited limited, cell assembly plasticity for multiple experiences, in agreement with prior findings regarding single linear track spatial experiences (*11, 37*). However, the P24-25 cuboid-reared rats had a markedly higher plasticity than the younger cuboid-reared rats, suggesting that as pattern separation for multiple spatial experiences improved with age, so did the ability to store these experiences. Interestingly, the deficits in pattern separation we observed in the P25 sphere-reared rats led to a marked reduction in the overall experience-induced plasticity in reactivation of cell assemblies in these rats (*16, 34–36*) during Post-Day2Runs sleep (multifold lower than in P24-25 cuboid-reared rats and comparable to that of younger P16-23 cuboid-reared rats; Fig. 6D). While the P25 sphere-reared rats had an impaired experience-dependent plasticity in reactivation, they exhibited a significant experience-dependent reorganization of cell assembly activity (i.e., higher in Pre-Day2Run1 sleep than Post-Day2Run2 sleep in some assemblies, higher in Post-Day2Run2 sleep than Pre-Day2Run1 sleep in others, indicative of an experience-dependent reorganization of the hippocampal network).

Based on these results, we propose that after sphere-rearing, the increased similarity of neuronal ensembles during sleep prevented the expression of unique preplay for distinct future run experiences, which contributed to reducing both pattern separation during track(s) exploration and plasticity in reactivation of those experiences.

### Experience with geometric linearity reconfigures neuronal ensembles during run largely countering the effects of sphere-rearing

We wondered whether the observed deficits induced by early-life deprivation from geometric linearity could be countered following experiences on multiple linear tracks across several days. Thus, on Day3 and Day4 we exposed our rats to alternating sessions of sleep and run on multiple linear tracks (Fig. 7A) and compared the changes in neuronal representations across days and groups. The tracks explored on Day3 were either previously experienced over multiple days (track 1, familiar) or were novel (tracks 4-5). This allowed us to study the influence of experience with geometric linearity on representations of previously experienced familiar tracks and, separately, the role of such experience in latently shaping hippocampal dynamics for representing novel tracks. Similarly, on Day4, we re-exposed our rats to a familiar track (track 4) followed by exposure to a novel track (track 6). Thus, our experimental protocol allowed the rats to accrue experience of geometric linearity across four days as they repeatedly experienced up to 6 distinct linear spatial contexts (tracks).

**Fig. 7.**
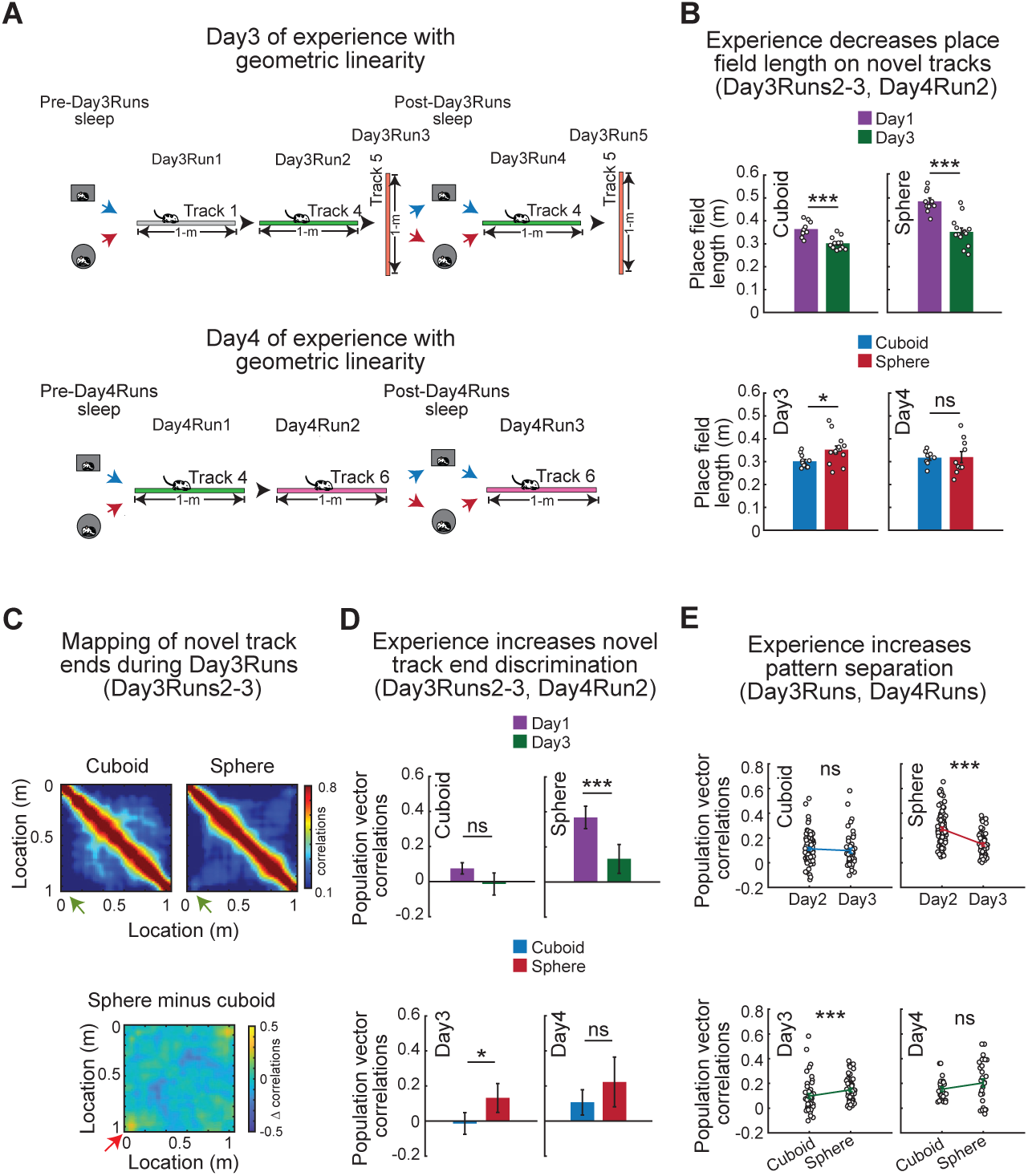
Repeated experience with geometric linearity across four days largely rescues the deficits observed during run associated with sphere-rearing from birth. (**A**) Experimental design on Day3 (top) and Day4 (bottom): alternating sleeps and runs on multiple linear tracks. (**B**) Experience-dependent changes in primary place field length on novel tracks across experimental days. Higher reduction in place field length across days in sphere-(top-right, p<0.0003) vs. cuboid-reared rats (top-left, p<0.0007). Significant residual difference between groups on Day3 (bottom-left, p<0.03), but not on Day4 (bottom-right, p>0.05, rank-sum tests). (**C**) Correlation matrix of neuronal ensemble mapping of locations along the linear track during Day3Runs2-3 for cuboid-(top-left), sphere-reared rats (top-right), and the difference between groups (bottom). Green/red arrows mark increased residual correlations on Day3 at track ends in sphere-reared rats. (**D**) Population vector correlation of place maps for track ends on novel tracks on Day3 compared with Day1. Note improved discrimination across days in sphere-(top-right, p<0.004) but not cuboid-reared rats (top-left, p>0.05) and the residual difference between groups on Day3 (bottom-left, p<0.03). Track end discrimination for novel tracks became similar for cuboid– and sphere-reared rats on Day 4 (bottom-right, p>0.05, rank-sum tests). (**E**) Increased pattern-separation of multiple tracks on Day3 over Day2 (top-right) in sphere-(p<10^-7^) but not cuboid-reared rats (top-left, p>0.05, rank-sum tests, left). Day3 pattern separation across groups: p<0.002, rank-sum test (bottom-left). Pattern separation of place cell maps for multiple tracks became similar across groups on Day4 (bottom-right, p>0.05, rank-sum test). N (rats/group x 2 directions/rat for each track) on Day3 and 4: Cuboid-reared (P25-26 and P26-27, respectively)=6; Sphere-reared (P26 and P27, respectively)=8. ***p<0.005. *p<0.05. ns=not significant.

We first examined the effect of experience on the properties of place cells and neuronal ensembles in the sphere-reared rats during runs on individual and across multiple linear spatial experiences (Figs. 1, 3 and 5). Experience over days with multiple linear tracks markedly improved spatial tuning properties of individual neurons. While primary place field length was reduced to a greater degree in sphere-reared rats for novel tracks on Day3 compared with Day1, it remained significantly higher in sphere-compared with cuboid-reared rats and became similar across groups on Day4 (Fig. 7B).

Mapping of novel linear space by neuronal ensembles of sphere-reared rats displayed less pronounced differences on Day3, when representations for the opposite track-ends during run were more distinct than on Day1 and became similar with those of cuboid-reared rats during Day4Runs (Fig. 7, C and D).

Finally, we investigated the effect that extended experience with geometric linearity had on the ability to pattern separate across multiple distinct linear tracks in cuboid– and sphere-reared rats. We observed an improvement in neuronal pattern separation for multiple linear experiences on Day3 compared with Day2 specifically in the sphere-reared rats (Fig. 7E), which became similar to that of the cuboid-reared rats by Day4 (Fig. 7E, bottom-right). Together, these results demonstrate that four days of experience with geometric linearity were sufficient to rescue the deficits in hippocampal neuronal representations of individual and multiple tracks after sphere-rearing.

### Experience with geometric linearity enriches the intrinsic sleep repertoire recruited during future experiences in sphere-reared rats

We next determined if and how experience with geometric linearity across days could reconfigure the intrinsic sleep network activity patterns as well as their selection by future experiences. The improvements in neuronal representation of linear space expressed during run in the sphere-reared rats on Day3 were preceded by expression of significant p/replay of future run trajectories during the preceding Pre-Day3Runs sleep sessions (all tracks explored within a run session following a sleep session, Fig. 8A, left), which became similar across groups by Day4 (Fig. 8A, right). Additionally, future track-specific p/replay emerged on Day3 in the sphere-reared rats (Fig. 8A, left) and became similar to that of cuboid-reared rats by Day4 (Fig. 8A, right).

**Fig. 8.**
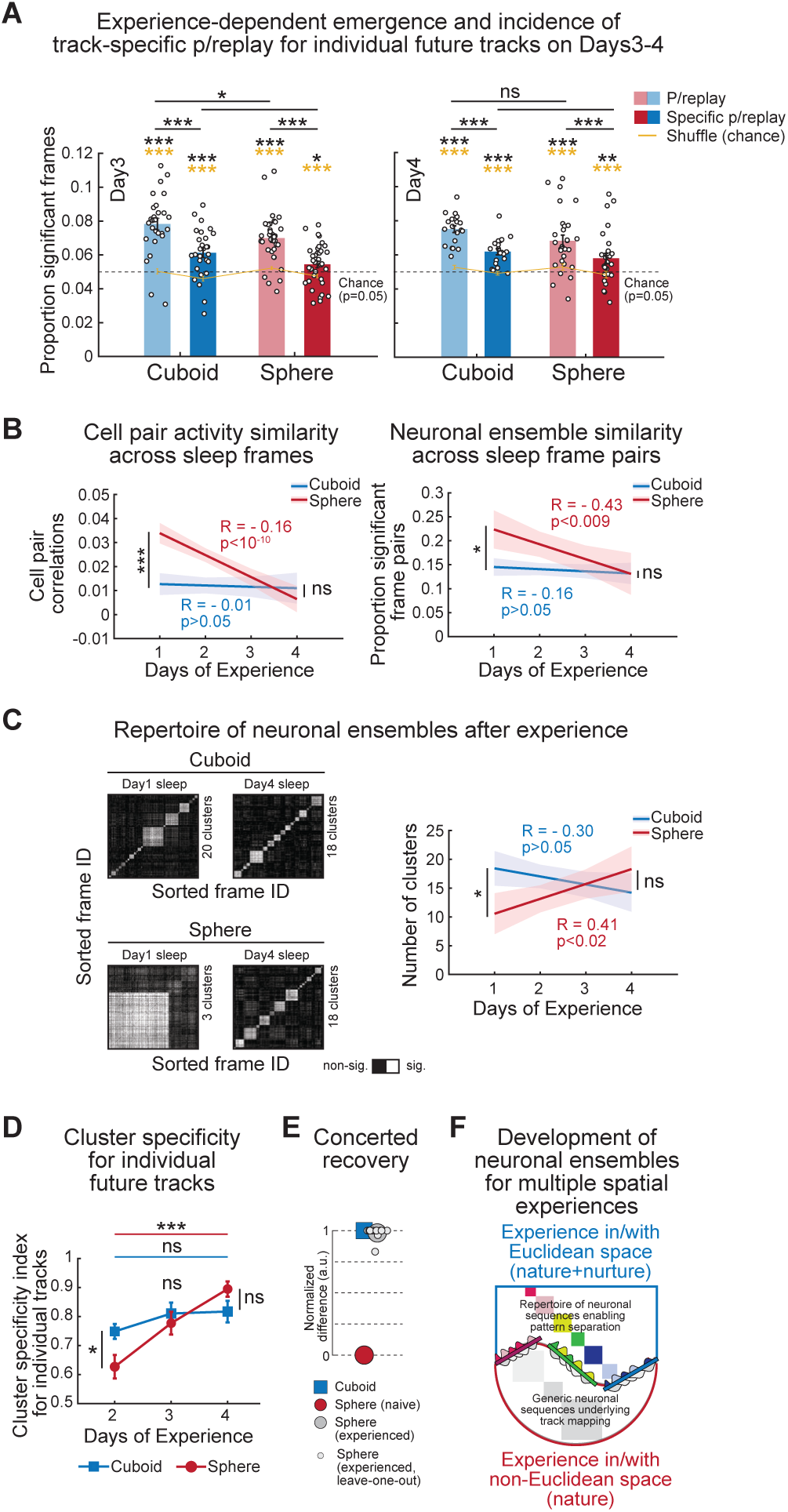
Repeated experience with geometric linearity across four days enriches the intrinsic sleep repertoire in sphere-reared rats. (**A**) Incidence of p/replay for future tracks is significantly above chance levels in both groups (left, cuboid vs. shuffle: p<10^-7^, sphere vs. shuffle: p<10^-7^, paired t-tests, yellow stars; p<10^-8^ t-test vs. p=0.05 chance for both groups, black stars; p<10^-10^, Binomial tests vs. p=0.05 chance for both groups). Incidence of track-specific-exclusive p/replay for future tracks is above chance levels in both groups (cuboid vs. shuffle: p<10^-4^, sphere vs. shuffle: p<0.002, paired t-tests, yellow stars; cuboid: p<10^-3^, sphere: p<0.02, t-tests vs. p=0.05 chance, black stars; p<10^-5^, Binomial tests vs. p=0.05 chance for both groups). P/replay and track-specific-exclusive p/replay remained higher in cuboid-reared rats on Day3 (p<0.05, t-tests, left). On Day4, p/replay (cuboid vs. shuffle: p<10^-8^, sphere vs. shuffle: p<10^-4^, paired t-tests, yellow stars; cuboid: p<10^-10^, sphere: p<10^-5^, t-tests vs. p=0.05 chance, black stars; p<10^-10^, Binomial tests vs. p=0.05 chance for both groups) and track-specific-exclusive p/replay (cuboid vs. shuffle: p<10^-5^, sphere vs. shuffle: p<0.005, paired t-tests, yellow stars; cuboid: p<10^-5^, sphere: p<0.006, t-tests vs. p=0.05 chance, black stars; p<10^-5^, Binomial tests vs. p=0.05 chance for both groups) for future tracks remained above chance levels and were similar in both groups (p>0.05, t-tests, right). (**B**) Experience-dependent reduction in the proportion of pyramidal cell pair correlations across sleep frames in sphere-reared rats (left, cuboid/sphere-reared Pearson’s correlations; Day1 sleep, sphere vs. cuboid, p<10^-10^; Day4 sleep, cuboid vs. sphere: p>0.05, rank-sum tests). Experience-dependent reduction in the proportion of significantly correlated neuronal ensembles across sleep frames in sphere-reared rats (right, cuboid/sphere-reared Pearson’s correlations; Day1 sleep, cuboid vs. sphere: p<0.03; Day4 sleep, cuboid vs. sphere: p>0.05, t-tests). (**C**) Experience on linear tracks increases the number of sleep-frame clusters exclusively in the sphere-reared rats (left, representative examples from same rats across days; right: cuboid/sphere-reared Pearson’s correlations; Day1 sleep, cuboid vs. sphere: p<0.006; Day4 sleep, cuboid vs. sphere: p>0.05, rank-sum tests). (**D**) Significant increase in cluster specificity for multiple individual tracks from Pre-Day2Run1 sleep to Pre-Day3Runs (p<0.02) and Pre-Day4Runs (p<10^-4^) sleep sessions in the sphere-but not cuboid-reared rats (p>0.05, t-tests), resulting in similar cluster specificity in sphere-vs. cuboid-reared rats on Pre-Day4Runs sleep sessions (p>0.05, t-test). (**E**) Concerted recovery of six individual neuronal parameters (see Methods for details) after 4 days of experience with geometric linearity in the sphere-reared rats. Recovery is quantified as proportion difference in parameter values between cuboid– and sphere-reared rats on Day4 (large grey dot) along a normalized arbitrary scale bounded by the corresponding values on Day1-2 in the sphere-(‘zero’; large red dot) and cuboid-reared (‘one’; large blue square) rats. Small grey dots reflect the contribution of each factor to the group average (large grey dot). (**F**) Cartoon model of the role of early-life experience in the development of the hippocampal neuronal ensemble repertoire. Colored/grey squares: frame-pair clusters (note the relative numbers across groups). Midline: 3 separate linear tracks with place cells recruited from color-matching clusters. N (rats/group x 2 directions/rat for each track) on Day3 and 4: Cuboid-reared (P25-26 and P26-27, respectively)=6; Sphere-reared (P26 and P27, respectively)=8. ***p<0.005. **p<0.01. *p<0.05. ns=not significant.

We next asked if the extended experience with geometric linearity influenced the intrinsic sleep dynamics as well (Fig. 4). We found that it reduced the similarity of pyramidal cell-pair activity across sleep frames (Fig. 8B, left) and the similarity of ensemble frame-pair activity during Day4 sleep sessions compared to Day1 sleep sessions in the sphere-reared rats (Fig. 8B, right). This increased experience with geometric linearity enriched the intrinsic repertoire of distinct neuronal ensembles during sleep increasing the number of distinct clusters, whose selection during run contributed to a better discrimination of the multiple, distinct linear track experiences in the sphere-reared rats (Fig. 8, C to D).

Altogether, repeated experience with geometric linearity over four days largely countered the multiple effects due to sphere-rearing from birth, as expressed during sleep, run, and in the relationship of sleep with run (Figs. 7 and 8; Fig. 8E models the concerted recovery of 6 major neural parameters). Intriguingly, these improvements in the representation of geometric linearity by the sphere-reared rats followed their highly controlled run experience on linear environments despite sleeping and resting in spherical sleep boxes and home cages during the day and overnight, respectively. These improvements might have been facilitated by our protocol, which involved repeated training on up to 6 distinct linear contexts with different geometric relationships. Overall, experience with geometric linearity at P24-27 after sphere-rearing from birth was sufficient to re/shape CA1 neuronal ensembles, which improved their distinct encoding and storage of multiple linear contexts.

## Discussion

We have demonstrated that depriving rats of experience with crucial features of Euclidean geometry via sphere-rearing across development affected individual place cell tuning for linear space. Yet sphere-rearing did not block the *emergence* of preconfigured and plastic hippocampal time-compressed neuronal sequences that encode and consolidate novel experiences on a linear track. Instead, early-life experience with linearity, vertical boundaries, planarity, and corners was critical for the emergence of an enriched repertoire of preconfigured neuronal sequences, which enabled discrimination of distinct spatial features and experiences on multiple linear tracks. Repeated experience with geometric linearity accrued over 4 days significantly reshaped neuronal ensembles, enriching their repertoire, and improved pattern separation of multiple experiences, largely reversing the initial effects of spatial deprivation. Thus, while dispensable for the representation of many features of a generic linear track experience (*38*), prior experience with Euclidean geometry proved more critical for the distinct representation of multiple linear track experiences and track ends and corners via rapid pattern separation during early development. From a nature vs. nurture debate perspective, early-life experience with features of Euclidean geometry (i.e., *nurture*) increases hippocampal network *performance* to rapidly discriminate between multiple linear environments, while network *competence* to express a pre-configured, generic representation of linear environments develops *a priori* (i.e., *nature*) (Fig. 8F).

Earlier studies did not discover a critical role of early-life sensory experience in the development of spatial circuits in the brain, including place cells and head-direction cells (*39, 40*), while a recent publication showed that early-life geometric experience can influence the expression of individual grid cells in the adult medial entorhinal cortex (*41*). Here we provided three lines of evidence for the role that specific early-life experience with spatial geometry had in sculpting p/re-configured hippocampal neuronal circuits though to be crucial for spatial representation and memory formation. First, geometric deprivation reduced both the intrinsic repertoire of preconfigured neuronal ensembles and their recruitment as distinct ensembles across multiple linear tracks, resulting in impaired plasticity for consolidating multiple experiences. Our findings indicate that a reduced neuronal ensemble repertoire observed in the pre-experience sleep in the naïve sphere-reared rats likely contributed towards the reduced network performance to discriminate multiple linear environments expressed as deficits in pattern separation. Notably, the network competence of sphere-reared rats to represent a sequential trajectory prior (e.g., preplay) and during (e.g., theta sequences) their very first encounter with a linear track developed *a priori*. This indicates that innate intrinsic developmental programs and/or experience with various aspects of the external world, other than Euclidean geometry, were sufficient to sustain the developmental emergence of time-compressed sequential patterns of neuronal firing previously assumed to require prior experience with geometric linearity. These findings suggest that crucial aspects of higher order ensemble representations of space-time emerge according to Kantian apriorism (*12*) rather than the predicaments of British empiricism (*42*). How intrinsic developmental programs and experiences with curved space enabled the emergence of network (pre)configuration expressed as preplay for future novel trajectories in linear environments (*9*) remains to be determined.

Second, rats’ extended experience with curved space inside the spheres likely contributed to the expression of changed (i.e., warped) and over-generalized representations for multiple distinct tracks and spatial features. In addition, linear borders, right angles, and vertical walls present in cuboid cages, but absent in sphere cages, may have acted as powerful geometric landmarks (*43, 44*) instructing brain networks to distinctively map start(s) and end(s) of multiple individual spatial trajectories (*32*). The absence of these geometric landmarks inside spheres deprived rats of this critical information. Consequently, the start and end of most trajectories taken by rats inside spheres (and distinct trajectories altogether) were likely to appear more similar, which may explain why the start/end portions of a linear track (and full distinct tracks) were mapped more similarly than in the cuboid-reared ones. During path integration, locations along rats’ trajectory on a linear track are mapped as a function of distance to track start in the earlier segments of the trajectory and distance to track end in the later segments (*45*). This integration/realignment process (as a form of distance coding) together with the more similar neuronal ensemble representation of the start and end of individual tracks might explain the changed representation of linear trajectories, which appeared as a warped representation of linear trajectories in the sphere-reared rats.

Finally, repeated experience with linear environments over four days largely reversed the effects of spatial deprivation by sphere-rearing. Interestingly, the effects of this spatial deprivation extended to new, not yet experienced linear contexts, indicating that early-life experience of geometry sculpts the neuronal ensemble rules for representing novel spaces. We hypothesize this rapid shift in neuronal representations of sphere-reared rats toward accurate depiction of geometric linearity was facilitated by our very selective deprivation from geometric linearity, which generally spared animal locomotion and age-dependent network maturation, and by the evolutionary and ecological affordances (*46*) of linear space for rats. Furthermore, the existence of preconfigured, preplay dynamics in the sphere-reared rats likely contributed to this shift as well. Whether and how the development of the (pre)configured hippocampal network repertoire for multiple spatial experiences occurred within a critical period early in life, similar to sensory brain networks (*47*), remains to be determined.

## Author Contributions

U.F. and G.D. collected and analyzed the data. G.D. conceived and designed the study. G.D. and U.F. wrote the manuscript.

## Acknowledgments

We thank K. Loetscher, B. Sanders and J. Sibille for help with data collection and analysis, and all the members of the Dragoi lab for comments and help. This work was supported by NIH grants R01NS10491, R01MH121372, and R35 NS132342 to G.D. The reported data are archived on file servers at the Yale Medical School.

## Competing interests

The authors declare no competing interests.

## Data and materials availability

All data needed to evaluate the conclusions in the paper are present in the paper and/or the Supplementary Materials.

